# How the level of reward awareness changes the computational and electrophysiological signatures of reinforcement learning

**DOI:** 10.1101/421743

**Authors:** C.M.C. Correa, S. Noorman, J. Jiang, S. Palminteri, M.X Cohen, M. Lebreton, S van Gaal

## Abstract

The extent to which subjective awareness influences reward processing, and thereby affects future decisions is currently largely unknown. In the present report, we investigated this question in a reinforcement-learning framework, combining perceptual masking, computational modeling and electroencephalographic recordings (human male and female participants). Our results indicate that degrading the visibility of the reward decreased -without completely obliterating- the ability of participants to learn from outcomes, but concurrently increased their tendency to repeat previous choices. We dissociated electrophysiological signatures evoked by the reward-based learning processes from those elicited by the reward-independent repetition of previous choices and showed that these neural activities were significantly modulated by reward visibility. Overall, this report sheds new light on the neural computations underlying reward-based learning and decision-making and highlights that awareness is beneficial for the trial-by-trial adjustment of decision-making strategies.

**Significance statement:** The notion of reward is strongly associated with subjective evaluation, related to conscious processes such as “pleasure”, “liking” and “wanting”. Here we show that degrading reward visibility in a reinforcement learning task decreases -without completely obliterating- the ability of participants to learn from outcomes, but concurrently increases subjects tendency to repeat previous choices. Electrophysiological recordings, in combination with computational modelling, show that neural activities were significantly modulated by reward visibility. Overall, we dissociate different neural computations underlying reward-based learning and decision-making, which highlights a beneficial role of reward awareness in adjusting decision-making strategies.

## Introduction

How we make decisions depends strongly on the outcomes that have been previously associated with the available courses of action. Actions that have been often linked with rewards (e.g. food, money) are more likely to be repeated than actions that have not been rewarded (or punished even) (Berridge & Robinson, 2003; Dayan & Balleine, 2002; Rangel, Camerer, & Montague, 2008). Generally, the notion of reward is strongly associated with subjective evaluation, related to conscious processes such as “pleasure”, “liking” and “wanting” (Berridge & Robinson, 2003). However, how human decision making changes depending on reward awareness is unclear. Assessing how the level of awareness of information changes or may bias value-based learning and decision-making may prove critical to understanding apparent irrationality observed in human behavior (Evans, 2008; Evans & Stanovich, 2013; Kahneman, 2003; Newell & Shanks, 2014; Weber & Johnson, 2009).

Rewards have two fundamental roles in the decision-making process. First, in decision situations, expected rewards act as incentives, which determine choices and increase the amount of motor or cognitive effort one is willing to expend to reach a goal (Berridge, 2004; Schmidt, Lebreton, Cléry-Melin, Daunizeau, & Pessiglione, 2012). Second, after a decision has been enacted and the action effectuated, the obtained reward -or absence of reward-drives important learning processes: successful actions are reinforced, while unsuccessful ones are discouraged (Sutton & Barto, 1998). Despite rewards being strongly associated with subjective feelings, notably to emotions and to the notion of expected pleasure (Berridge & Robinson, 2003), recent studies have reported that reward cues that are masked from awareness can still directly influence task performance (Aarts et al., 2008; Bijleveld, Custers, & Aarts, 2012; Capa, Bouquet, Dreher, & Dufour, 2013; Pessiglione et al., 2007). These results suggest that the first role of reward information –incentivizing decision and effort production-may be processed outside the scope of awareness in the human brain to facilitate human performance (but see Bijleveld et al. 2014 for results challenging this view). On the other hand, little is known about if and how the second role of rewards –i.e. the propensity to *reinforce* successful actions - is modulated by awareness.

To address this question, thirty-two participants performed a probabilistic reversal learning task in which we manipulated the visibility of reward using a standard masking technique. Participants were instructed to choose one of two response options, which led probabilistically either to a significant reward (50 cent coin, “reward condition”) or a negligible one (1 cent coin, “no-reward condition”). Response-reward contingencies reversed several times over the course of the experiment and participants were instructed to select the response option that was most often rewarded (**Fig. 1a**). Masked and unmasked feedback were mixed within blocks to explore the relative weighting of both types of feedback. We combined EEG measurements with computational modeling to investigate, at the time of reward processing and on a trial-by-trial basis, the neural correlate of the different processes influencing participants’ future choices and how those were affected by reward visibility. Thereby, the present work builds on previous studies that have linked reinforcement learning (RL) models to human neural data obtained from both fMRI and EEG measurements (Daw, Gershman, Seymour, Dayan, & Dolan, 2011b; Debener et al., 2005; Fischer & Ullsperger, 2013; Fouragnan, Queirazza, Retzler, Mullinger, & Philiastides, 2017; Hauser et al., 2014; O’Doherty et al., 2007; Ullsperger, Fischer, Nigbur, & Endrass, 2014). In line with previous work, event-related potential (ERP) analyses focus on the feedback-related negativity (FRN) and the P3 component (Holroyd, Nieuwenhuis, Yeung and Cohen, 2003; Holroyd & Coles, 2002). The investigations on the EEG-correlates of RL learning concentrate on three (computational) variables: the prediction-error (signed PE), the level of surprise (unsigned PE) and switch/stay behavior on the next trial (Cohen & Ranganath, 2007; Collins & Frank, 2018; Fischer & Ullsperger, 2013; Fouragnan et al., 2017). This approach allows us to investigate the impact of reward visibility on different cognitive processes involved in probabilistic reward-guided learning.

**Fig. 1.**
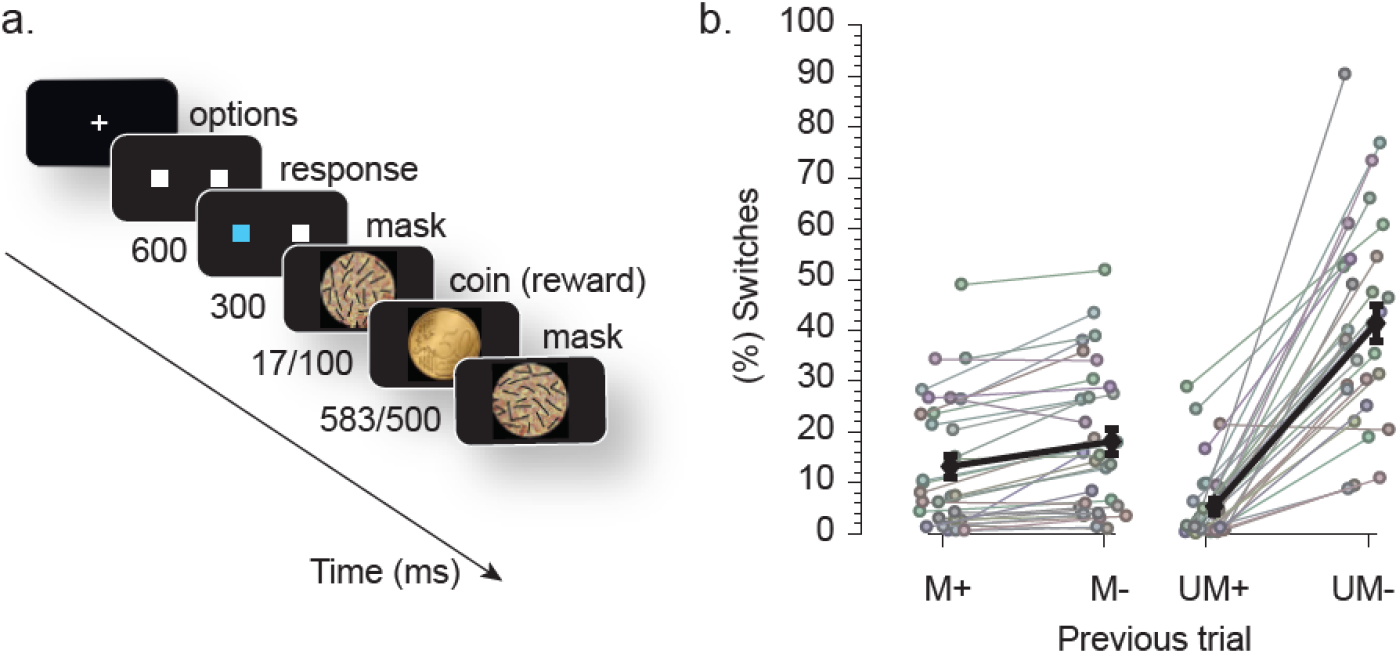
Experimental setup and behavior. **(a)** Two response options (white boxes on the left/right of fixation) were shown on the screen until a response was given. A correct response was rewarded with a 70% probability (50 cent coin) and not rewarded with a 30% probability (1 cent coin). Reward visibility was manipulated by masking. Unmasked (long coin presentation, short backward mask presentation) and masked (short coin presentation, long backward mask presentation) reward trials were mixed within blocks and randomly chosen across trials (each with a 50% probability). Which response option was most rewarded changed every 75-125 trials. **(b)** The percentage of switches, at the group level (in black) and for individual subjects (in gray) after specific trials. M: masked; UM: unmasked; +: reward; -: no-reward; error bars represent ±s.e.m.

## Materials and methods

### Participants

Thirty-two students from the University of Amsterdam (8 males, 24 females, aged 22.25±3.1) participated in the experiment for course credits or financial compensation. All participants gave their written informed consent prior to participation, had normal or corrected-to-normal vision and were naive to the purpose of the experiments. All procedures were executed in compliance with relevant laws and institutional guidelines and were approved by the local ethical committee of the University of Amsterdam.

### Task

Stimuli were presented using Presentation software (Neurobehavioral Systems, Inc) against a black background at the center of a 20-inch VGA monitor (frequency 60 Hz), which was viewed by the participants from a distance of approximately 80 cm. Participants should fixate at the center of the screen and choose between a left or a right box distant 15 cm from each other by pressing a correspondent left or right chair button (parallel button).The chosen square was illuminated in blue for 600 ms, indicating the participants’ response followed by a reward (50 cent coin) or a punishment (1 cent coin) that could be shown on a visible (100 ms) or masked (17 ms) way. Stimuli were used similarly to those by (Zedelius, Veling, & Aarts, 2012). A variable time, 1500 to 2500 ms inter-trial-interval separated each trial. If participants did not select a target after 1500 ms, a “too late!” message was displayed (see **Fig. 1a**).

Sides were rewarded in a 70/30% fashion. This probability condition was reversed several times along the 1200 trials so that, in order to decide advantageously, participants had to keep track of eventual “rule changes”. We refer to the choices made on the 30% probability side as ‘incorrect’ choices, and those made according to the 70% rewarded side as “correct” ones. Probabilities were fixed across trials within blocks, which lasted 75–125 trials. The block length had a minimum value, but it was dependent on how fast participants could learn the rule at stake. In order to assure that everyone could learn the probabilities, for at least 10 trials in a row they should have been able to choose the “correct side” option for more than 60% of the last 25 trials, otherwise additional trials could be added until this condition was completed. Self-paced rest breaks were given every 70 trials, presenting participants the percentage of correct sides they have chosen according to the rule at stake. This break never coincided with changing probabilities conditions and participants were told about that.

In 10% of the trials a forced choice discrimination question asked “Which coin did you just see?” while displaying a 1 cent or a 50 cent coin. This question was randomly asked for the same amount of times for visible and masked coins. Participants were instructed that the probability of the correct response being a 1 cent or 50 cent coin was 50%. Participants were explained that they would get paid according to their performance at the end of the experiment. Finally, all participants received a bonus of €5 on top of what they had already received. Participants were instructed to choose one of the two targets on each trial, to pay attention to the reward, and to try to win as much money as possible.

### Models Building Blocks

We designed 18 different models, all adapted from a Q-learning model. Our Q-learning included 3 basic modules: learning, choice and perseveration (see **Fig. 2a**).

**Fig. 2.**
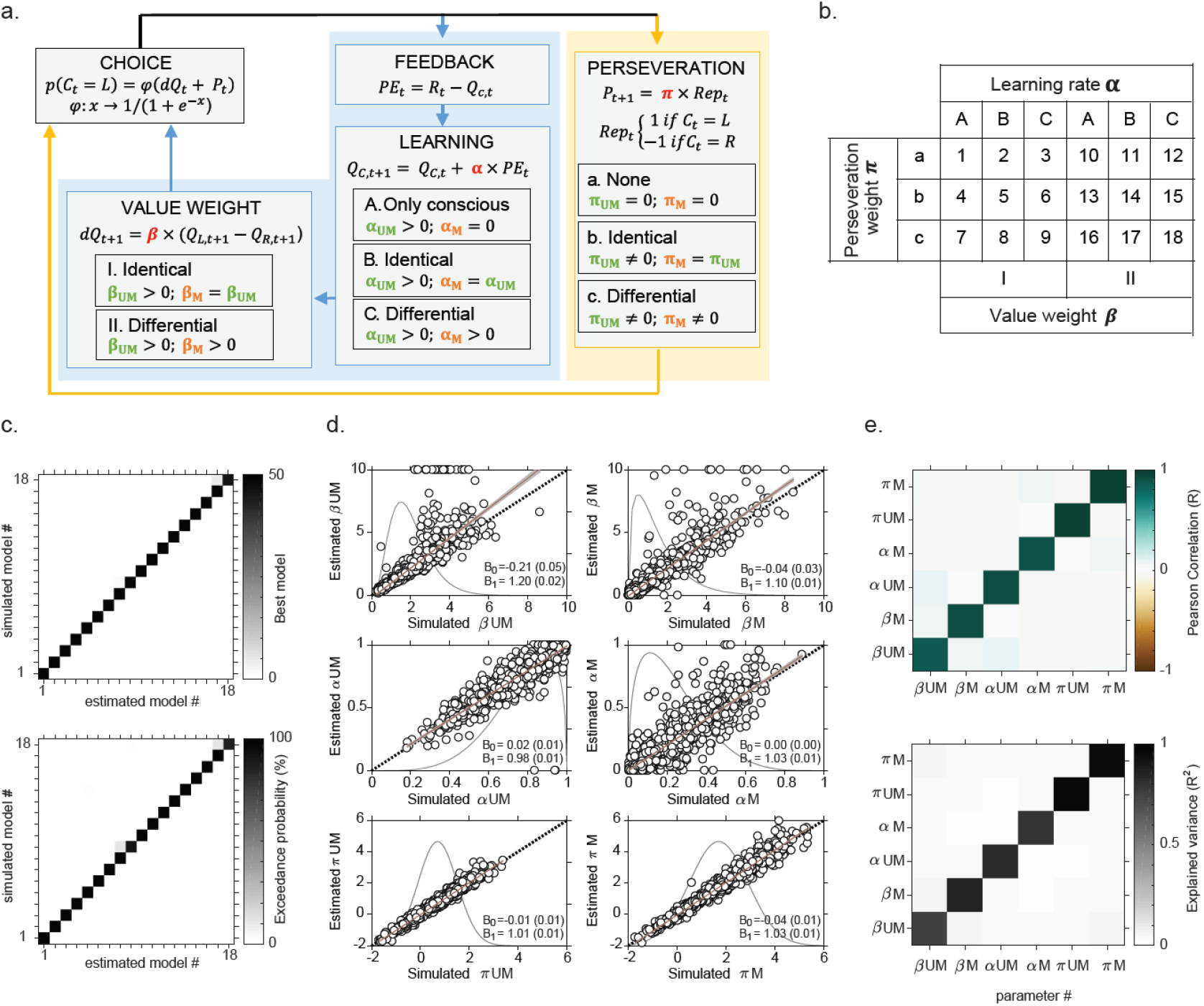
Modeling approach. **(a)** The computational architecture used to build the model space**. (b)** Model space. Eighteen models were built by systematically combining the different options available for the different computational modules. **(c) Model identifiability analysis**. Data from 32 synthetic participants were simulated with each of our 18 models. Bayesian model selection was used to identify the most probable model generating the data, using model exceedance probability. This procedure was repeated 50 times. Overall, all 18 models were correctly identified more than 90% of the time (>45 out of 50 simulations, see top confusion matrix), with an average exceedance probability > 90% (bottom confusion matrix). **(d) Parameter recovery analysis - general**. Overall, data from 1600 synthetic participants (50 simulations x 32 individuals) were simulated with the full model (model 18). The 6 estimated parameters per participants were then regressed against the true parameters used for simulating the data. Results show very good identifiability, with regression intercepts (β_0_s) close to 0, regression slopes (β_1_s) close to 1 and highly significant (all p-values lower than Matlab’s precision –i.e. reported as = 0). Each dot represents a synthetic individual. The black dotted lines represent the identity line, the red continuous lines the best linear fits, and the shaded grey areas the 95% confidence interval around the best-linear fit. The grey densities represent the probability distributions used to sample the parameters. **(e) Parameter recovery analysis – individual simulations**. The confusion matrices represent summary statistics of the correlations between parameters, estimated over 32-subjects simulations, and averaged over the 50 simulations. Diagonal: correlations between simulated and estimated parameters. Off diagonal: cross correlation between estimated parameters. Top: Pearson correlation (R). Bottom: explained variance (R^2^).

**Learning**. The basic idea is that participants learn by trial and error to compute a value Q for each option (choosing the left or the right cue). At each trial *t*, after a choice is made and the outcome of the choice *R*_*t*_ is revealed, the Q-value of the chosen option (Q_*C,t*+1_) is updated by integrating a so-called prediction-error δ_*t*_, which compares what was expected (Q_*C,t*_) to the actual outcome:

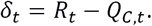

This update is typically scaled by a learning rate α, such that:

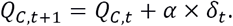

**Choice**. To account for the fact that people try to maximize their expected outcome, but can make errors or explore locally sub-optimal options, the choice (*C*_*t*_) is typically implemented as a softmax function:

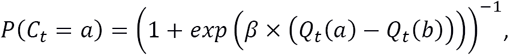

where *β* is the slope of the logistic choice function –the inverse temperature parameter-which we refer to as the value weight.

**Perseveration**. In order to capture the tendency of participants to stick to their previous choices independently of the received reward, we also included a perseveration bias π_*t*_ in the choice function. This function becomes:

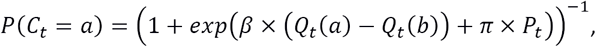

where

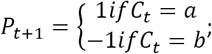

and *π* governs the weight of the past choice on the present decision –referred to as the perseveration weight.

When both learning and perseveration are present, the relative importance of *β* and *π* allow the model capture participants tendency to trade-off between sampling from learned value (*β*) vs simply repeating previous choices (π).

*Model Space*

Given that our task incorporates two types of reward - masked vs. unmasked - several scenarios are possible for learning and perseveration, which can be accounted for by different models. We first assumed that all models share a common basic block, that is, people learn from unmasked reward. Additionally, people can learn from masked reward, either at the same pace or at a different pace than after unmasked reward. Likewise, the value weight parameter can be identical or different after unmasked vs masked reward. As for the perseveration, it can be absent after both masked and unmasked reward, present and of identical strength, or present with different strengths. Those 3 learning, 2 choice-temperature and 3 perseveration scenarios were therefore combined, generating 18 possible models in our model space (**Fig. 2a/b**).

### Parameter optimization

We optimized the models free-parameters (α’s, β’s and π’s) by minimizing the negative log likelihood (LLmax) of the participant observed choices under the model using Matlab’s fmincon function, initialized at multiple starting points of the parameter space.

### Model Comparison

Negative log-likelihoods (LLmax) were used to compute the Bayesian information criterion (BIC), for each model, at the individual level (*BIC* = 2 × (*LLmax*) + *df* × log(n_*trial*_)), and used to approximate the model evidence (*e* = -*BIC*/2). Individual model evidences were then fed to the mbb-vb-toolbox (https://code.google.com/p/mbb-vb-toolbox/) to run a Bayesian Model Comparison (Daunizeau, Adam, & Rigoux, 2014). This Bayesian procedure estimates, among other criteria, the exceedance probability (denoted XP) for each model within a set of models, given the data gathered from all participants. Exceedance probability quantifies the belief that the model is more likely than all the other models of the set. An exceedance probability greater than 95% for one model within a set is therefore typically considered as significant evidence in favor of this model being the most likely. In addition, the relative BIC (δBIC, i.e., the BIC for each model relative to best model), can be used to compare models based on the Bayes factor scale proposed by Kass and Raftery (Kass & Raftery, 1995).

### Model identifiability and parameter recovery

We ran 50 simulations, generating choice patterns for cohorts of 32 synthetic subjects with the 18 different models in our model set. For those simulations, parameters were randomly sampled from probability distributions which approximate the distribution of parameters estimated from fitting the complete model (i.e. model 18) to the choices of our 32 participants. As is common in the field (Daw et al., 2011; Palminteri, Khamassi, Joffily, & Coricelli, 2015), inverse temperature parameters were sampled in Gamma distributions defined by a shape (a) and a scale (b) parameter (unmasked: a = 4.0; b = 0.5; masked: a = 1.5; b = 1.0) and learning rates were sampled in Beta distributions defined by 2 parameters α and β (unmasked: α = 5.0; β = 1.5; masked: α = 1.5; β = 5.0). Finally, perseveration parameters were sampled in Normal distributions, characterized by mean (μ) and standard deviation (σ) (unmasked: μ = 0.7; σ = 0.8; masked: μ = 1.7; σ = 1.2). Task properties and contingencies (block lengths, etc.) used for the simulations were rigorously identical to the 32 instances that participants faced in our experiment.

Then, we ran our Bayesian model-comparison (BMC) analysis on those 50×18 different simulations, and checked that all models are identifiable, i.e. can be correctly estimated as the most probable model in the set of 18 models by the BMC approach when they were actually used to generate the data. This first analysis intends to verify that nothing in the design of the model set, the parameter estimation or the model comparison approach, unduly advantages model 18 (e.g. that it is the most complex model), leading to mistakenly over-estimate the probability that model 18 explains our participants’ choices *in lieu of* other models. Next, because our models are nested, we assessed the parameter recovery in the full-model case (model 18): we computed the Pearson correlation between the parameters used to generate the data, and the parameters estimated by the maximum-likelihood fitting procedure. Additionally, we estimated the correlation between estimated parameters.

### Parameters and model recovery

All 18 models are correctly identified more than 90% of the time, with an average exceedance probability > 90% (**Fig. 2c**). A closer look at the parameters estimated from the 1200 subjects over the 50 simulations ran with model #18 (the most complex model, in which all other model are nested) show that parameters are also very well recovered, with regression intercepts (β_0_s) close to 0, regression slopes (β_1_s) close to 1 and highly significant (all p-values lower than Matlab’s precision –i.e. reported as = 0) (**Fig. 2d**). At the scale of a single simulation, the correlation between simulated and estimated parameters over 32 synthetic participants was very significant (averaged Pearson correlation = 0.92 and averaged R^2^= 0.85; **Fig. 2e**; diagonals), while no cross-correlation was observed between parameters (all R^2^< 0.06; **Fig. 2e**; off-diagonals).

### EEG measurements

EEG data was recorded and sampled at 512 Hz using a BioSemi ActiveTwo system. Sixty four scalp electrodes were measured, as well as 4 electrodes for horizontal and vertical eye-movements (each referenced to their counterpart) and 2 reference electrodes on the ear lobes (the average was used for referencing). After acquisition, standard pre-processing steps were performed in EEGLAB toolbox in Matlab. Data were bandpass filtered from 0.5 to 40 Hz off-line for ERP analyses. Epochs ranging from 1.8 s before to 2 s after reward presentation were extracted. Linear baseline correction was applied to these epochs using a -200 to 0 ms window. The resulting trials were visually inspected and those containing artifacts were removed manually. Moreover, electrodes that consistently contained artifacts were interpolated. Finally, using independent component analysis, artifacts caused by blinks and other events not related to brain activity were removed from the EEG data.

### ERP analyses

We focused on ERP components related to reward outcome processing with different latencies and topographical distributions. To zoom in on these specific components a central region of interest (ROI) was defined comprising of 15 midline electrodes (Fz, F1, F2, FC1, FCz, FC2, Cz, C1, C2, CPz, CP1, CP2, Pz, P1, P2), where both the relevant components can be observed (fronto-central FRN and ventro-parietal P3) (Chase, Swainson, Durham, Benham, & Cools, 2011; Cohen, Elger, & Ranganath, 2007; Cohen, Wilmes, & van de Vijver, 2011; Ullsperger et al., 2014a). Selecting a predefined ROI limits the number of comparisons that need to be performed, but we note that the results were robust and were not dependent on the specific sets of electrodes used as a ROI (Fig. 4). We investigated the effect of reward outcome separately for masked and unmasked trials. To correct for multiple comparisons due to the number of time-points tested, p values were FDR-corrected at an alpha-level of 0.05. All statistical analyses were performed in Matlab (Mathworks). Based on this ERP analysis three time-windows of interest were selected for follow-up analyses in which we related model parameters to single trial EEG responses.

**Fig. 4.**
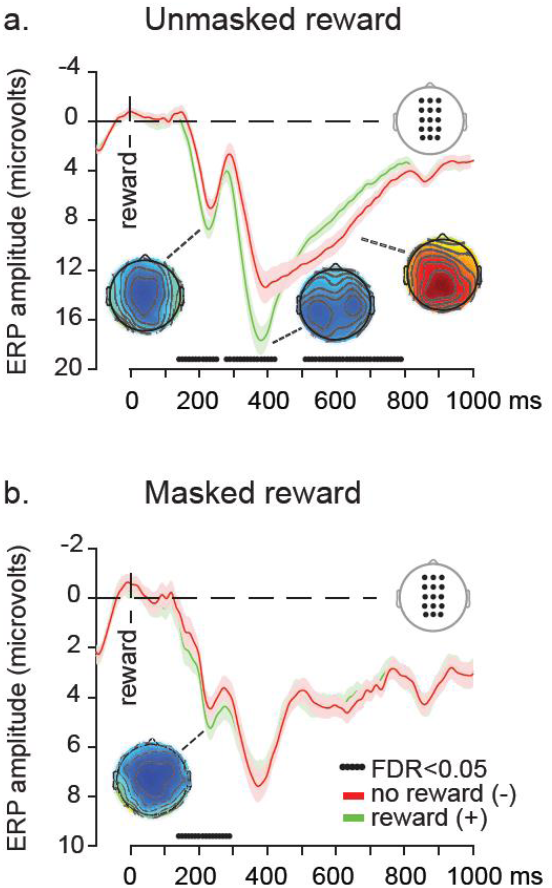
ERP results. ERPs for no-reward (red lines) and reward (green lines) for unmasked **(a)** and masked conditions **(b)**. Time=0 ms is reward presentation. The lower dotted black lines indicate significant time-windows, FDR corrected across the entire ERP time-window (p<0.05). Topographical distribution maps of the reward valence effect (no-reward minus reward, - vs +) were taken from the three broad time-windows (100-300 ms, 300-500 ms and 500-800 ms; scaling maps unmasked reward from left to right: [- 2:2], [-5:5], [-2:2]; scaling maps masked reward: [-2:2]). Error bars represent ± s.e.m.

### Single trial regression analyses

Multiple regressions of ERP amplitude on three model parameters were conducted. For each subject, each electrode, and each time point, the three parameters (PE, |PE|, switch/repeat on the next trial) were entered as predictor variables, and the ERP amplitudes as observations in the regression model. We checked that the correlations between the time-series of the 3 predictors was low (absolute value of pearson’s R averaged over subjects <.2), resulting in low multi-collinearity indices (variance inflation factors: VIF_PE_= 1.0596±0.0099; VIF_|PE|_= 1.0524±0.0147; VIF_switch/repeat_= 1.0712±0.0145). Beta-coefficients assigned to each predictor column, which reflect the regression weights between each of the three parameters and ERP amplitude, were estimated at the individual level, separately for each electrode and time point. The significance of the predictors was assessed at the population-level using random-effects (t-tests) on the regression coefficients averaged across the predefined time windows (100-300 ms, 300-500 ms, 500-800 ms) and the predefined ROI.

### Code availability

The codes used to analyze data from the current study are available from the corresponding author on reasonable request.

### Data availability

The datasets generated during and/or analysed during the current study are available from the corresponding author on reasonable request.

## Results

### Behavior

Participants were able to perform the task well and they accurately tracked probability reversals (mean correct response = 71.3±1.51%). In order to assess the reward discriminability in the masked (M) and unmasked conditions (UM), we computed participants’ d-prime, an unbiased measure of stimulus visibility, from the forced-choice discrimination trials that were presented throughout the task (10% of all trials, hence 120 trials in total). Although the overall discriminability was low in the masked condition, both masked and unmasked conditions exhibited above-chance accuracy in this discrimination test (UM: 96±1.15% correct, d’=3.97±0.14; t_31_=28.38, p<0.001; M: 55.7±1.13% correct, d’=0.35±0.07; t_31_=4.91, p<0.001). Given that chance-level performance on such a forced-choice discrimination task is a typical criterion used to show that participants are unable to perceive a stimulus consciously (Overgaard & Sandberg, 2012; Sandberg, Timmermans, Overgaard, & Cleeremans, 2010), this result implies that we cannot consider that the masked reward was nonconscious in all participants and for all trials.

Having established that participants performed the task correctly, we turned to a typical behavioral analysis of learning. Following previous studies (Chase et al., 2011; den Ouden et al., 2013), we computed participants switch rates after positive and negative outcomes, in both unmasked and masked conditions. Critically, participants switched their response more often after no-reward than after reward, and did so in both the unmasked and in the masked condition (unmasked: difference 36.06±0.59%, t_31_=10.76, p<0.001; masked: difference 4.90±0.15%, t_31_=5.65, p<0.001). The fact that participants tended to switch their choices significantly more after no-reward (1 cent) versus reward (50 cent) is generally interpreted as evidence for learning. It would therefore be tempting to conclude that our participants significantly learned from both unmasked and masked reward. However, this interpretation of switch patterns may not be devoid of statistical confounds, especially in designs where conditions (in this case masked and unmasked) are intermixed. Indeed, this pattern of results could easily be produced by participants learning the value of options from unmasked rewards and deriving all choices from those values - i.e. in the total absence of learning from masked reward. This is why we turned to model-based behavioral analyses that are devoid of this statistical confound, aiming at showing that learning from masked reward outcomes is still present when these issues are taken into account.

### Computational modeling

A simple delta-rule was used to capture how individuals updated the value of the chosen options after receiving reward. Following classical associative learning algorithms, the extent to which previous reward is integrated in the future option value was controlled by a *learning rate* α. Choices were derived from a logistic (soft-max) choice function, on the difference between option values. The slope of this choice function – typically referred to as choice temperature - was defined as *the value weight* β. Although very popular and accounting for a wide range of behavior, this learning mechanism might not account for the full choice pattern of participants in our task: indeed, within blocks, our participants might identify the best option and therefore start disregarding the feedback, putting more weights on their priors. To account for this behavior, we added a perseveration module to our computational model. Perseveration – defined as the tendency to repeat a choice regardless of the previous outcome - was integrated as an additional “bias” in the choice function, which regulated the probability of choosing the same option as that in the previous trial (den Ouden et al., 2013; Rutledge et al., 2009; Seymour, Daw, Roiser, Dayan, & Dolan, 2012; Voon et al., 2015). The extent to which perseveration contributed to the final choice was determined by a *perseveration weight* π (see **Fig. 2a**, **Materials and Methods**). We then systematically explored how masked versus unmasked reward impacted those different modules, by creating sets of models allowing – or not allowing - parameters to differ between those two conditions (see **Materials and Methods** and **Fig. 2b**). We thereby built 18 different models, which were subsequently fit to the behavior, using a maximum likelihood procedure. A model recovery (**Fig. 2c**) and a parameter recovery (**Fig. 2d-e**) analysis confirmed that our modelling approach is suitable to address our questions of interests (Palminteri, Wyart, & Koechlin, 2017) (see also **Materials and Methods**).

Regarding our participants’ data, a Bayesian model comparison approach identified model 18 as the best among our designs to explain the behavior (exceedance probability>80%, see **Fig. 2c**). The best fitting model differentiates *learning rate, value weight*, and *perseveration weight* parameters after unmasked and masked reward. Importantly, because our model space included models explicitly omitting learning from masked reward (**Fig. 2b**), this model comparison result demonstrates the existence of learning from masked reward, even when perseveration effects are taken into account.

Participant-level data reveals that the best fitting model gives a very good account of participant’s learning and switch behavior (average likelihood per trial=78.70±2.11%; **Fig. 3a** for three representative participants: s10, s20, s30). We then turned to the analysis of the best fitting model parameters (**Fig. 3b**). Learning rates appeared to be higher after unmasked than masked reward (α_UM_=0.67±0.03; α_M_=0.19±0.02, t_31_=17.01, p<.001), and so did value weights (β_UM_=1.94±0.18; β_M_=0.93±0.12, t_31_=7.24, p<0.001). However, the opposite was found for the weight put on previous choices (π_UM_=0.67±0.15; π_M_=1.67±0.21, t_31_=-4.72, p<0.001) (**Fig. 3b**).

**Fig. 3.**
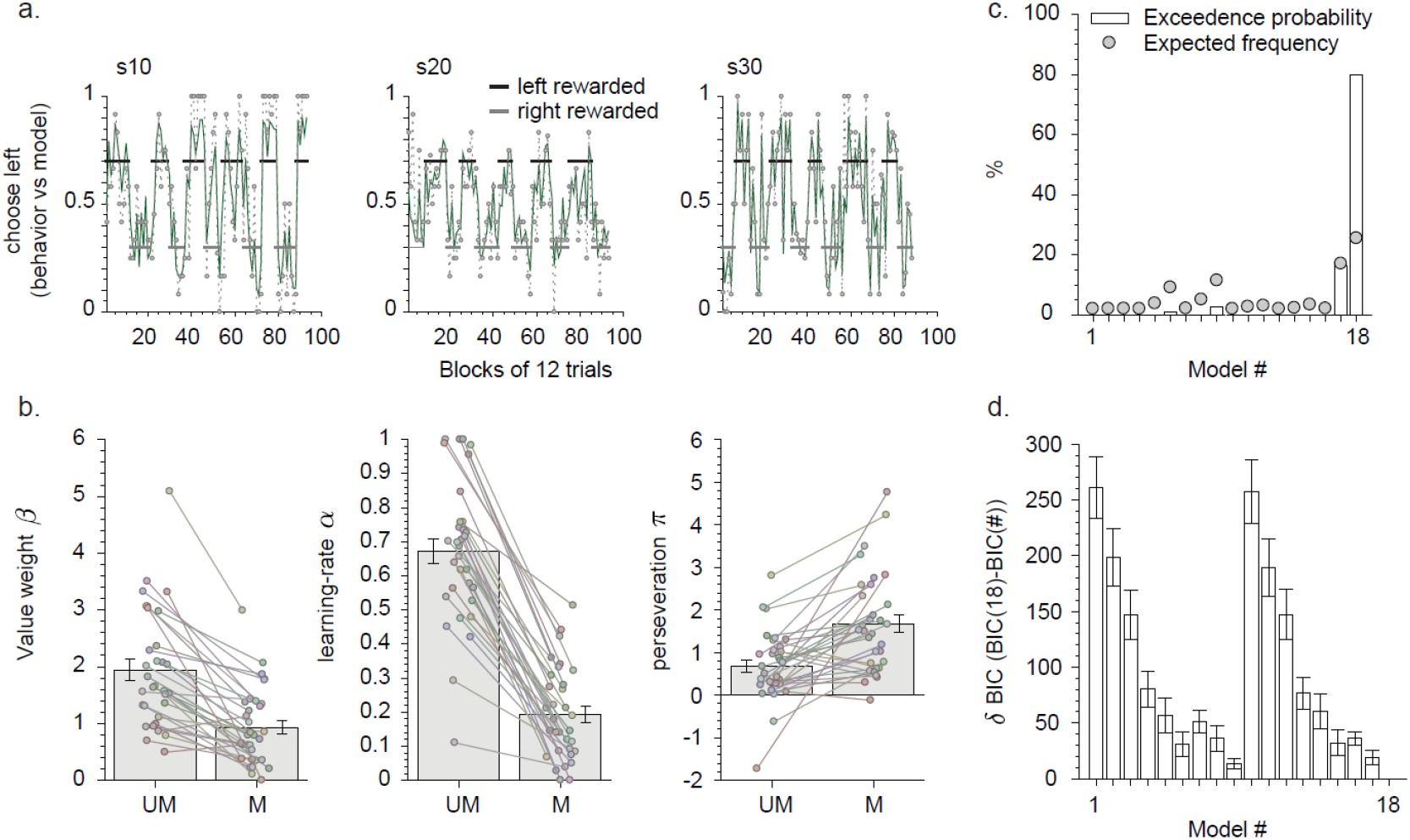
**(a)** Time course of the learning task by three representative participants (participant numbers 10, 20 and 30). The x-axis represents blocks of trials during the experiment and the y-axis represents the local fraction of left-hand responses selected by the participant. Thick black and gray lines represent the reward probability in the different blocks (75-125 trials). Gray-dotted lines represent the local fraction of left-hand responses. Green thick line represent the local probability of left-hand responses predicted by the computational model. Both behavioral choices and model predictions are averaged over 12 trials bins, and aligned on block transitions. **(b)** Model parameters for masked and unmasked conditions. Left: value weight. Middle: learning rate. Right: perseveration weight. M: masked reward, UM: unmasked reward. Histograms and error bars represent mean ± s.e.m. Connected dots represent individual parameters. **(c)** Model comparison. Results of a Bayesian model comparison analysis on our participants’ data. White histograms indicate the exceedance probability of each model, and grey dots their expected frequencies. **(d)** Relative BIC. Bayesian Information Criterion (BIC) of each model, compared to the best fitting model BIC (model 18). BICs are computed at the individual level (random effects). Histogram and error bars represent mean ± s.e.m.

These results lead to several crucial insights concerning reward learning. First, they demonstrate the existence of robust learning from masked rewards. Second, they clearly illustrate changes, due to reward visibility, in the trade-off between the tendency to base choices on the learned options’ values, and the tendency to repeat previous choices regardless of previous outcome. This thus suggests that the reliance on the longer term priors, based on the accumulation of recent choices, is increased when the outcome on the current trial is masked and therefore unreliable.

Finally, we ran independent linear regressions with each of the individual parameters from the model (6 parameters in total) as independent variables and overall performance (percentage correct) as dependent variable to explore what model parameters correlate with individual performance. Results show that inverse temperatures (β_UM_: beta=0.060, p<0.001; β_M_: beta=0.076, p<0.001) and perseveration parameters (π_UM_: beta=0.043, p=0.016; π_M_: beta=0.046, p<0.001) are positively correlated with performance, while learning rates (α_UM_: beta=-0.210, p=0.0016; α_M_: beta=-0.229, p=0.036) are negatively correlated with performance.

### ERPs and model-based EEG results

Having established, thanks to the manipulation of reward visibility, a clear computational dissociation between the contributions of learning versus choice perseveration to the behavior of our participants, we next aimed at dissociating the neural signatures of those components by leveraging electrophysiological recordings. In order to first identify the electrophysiological time-windows of interest, we performed an ERP analysis of reward-related activity, contrasting reward versus no-reward outcomes, at our central region of interest, which was based on previous studies (Cavanagh, Frank, Klein, & Allen, 2010; Cohen et al., 2011; Ullsperger et al., 2014) (see **Materials and Methods**).

Our analysis of event-related potentials revealed three significant events in the neural signal evoked by fully conscious (unmasked) outcomes: an early Feedback-Related Negativity (FRN) at fronto-central electrodes (“early” event), which was followed by a second, more centrally distributed negative component (“middle” event), and a final parietal P3 component (“late” event) (**Fig. 4a**, FDR corrected across time, p<0.05). Crucially, while masked outcomes also elicited an early fronto-central FRN, neither the second negative ERP component nor the P3 component could be observed in the masked condition (FDR corrected across time, p<0.05, **Fig. 4b**).

In order to relate the contributions of the different computational modules identified in our best fitting model (model 18, **Fig. 2**) to electrophysiological signatures of outcome-guided decision-making, we then turned to a model-based analysis of the EEG signal. In each participant, at each electrode and at each time point, we estimated a multiple regression with the trial-wise time-series of electrophysiological activity as the dependent variable, and trial-wise time-series of latent variables as independent variables (see **Experimental Procedures**). Three such independent variables, derived from our best fitting model, were included in this multiple regression: the signed prediction error, the unsigned prediction-error (typically interpreted as a measure of surprise (Cavanagh & Frank, 2014; Pearce & Hall, 1980)), and a variable indexing whether participants switched or repeated their choice from the previous to the next trial, which is directly related to the perseveration process (switch/stay behavior). Previous research has shown the existence of temporally overlapping but spatially separate contributions of the signed prediction error, reflecting the valence of the prediction error (positive or negative) and the unsigned prediction error (the absolute degree of expectation violation also referred to as surprise) to reward learning (Fouragnan et al., 2017).

In our model-based analyses, we focus on the three contiguous time-windows in which the model-free effects were most pronounced (early: 100-300 ms, middle: 300-500 ms and late: 500-800 ms). The signed PE regression results showed two clear peaks strongly overlapping in time with the early two ERP components that were revealed in the model-free ERP analysis (**Fig. 5a**). For both masking conditions, the signed prediction error was encoded in the early FRN (unmasked: t_31_=6.8, p<0.001; masked t_31_=4.2, p<0.001, difference: t_31_=3.0, p=0.005, early time-window). Similar results were obtained for the mid-latency negativity (unmasked: t_31_=11.2, p<0.001; masked: t_31_=3.0, p=0.005; difference: t_31_=8.1, p<0.001, middle time-window). In contrast, the later P3 component appeared only reached significance in the masked outcome conditions, although both conditions did not differ significantly (unmasked: t_31_=0.85, p=0.40; masked: t_31_=4.1, p<0.001, late time-window, **Fig. 5a)**.

**Fig. 5.**
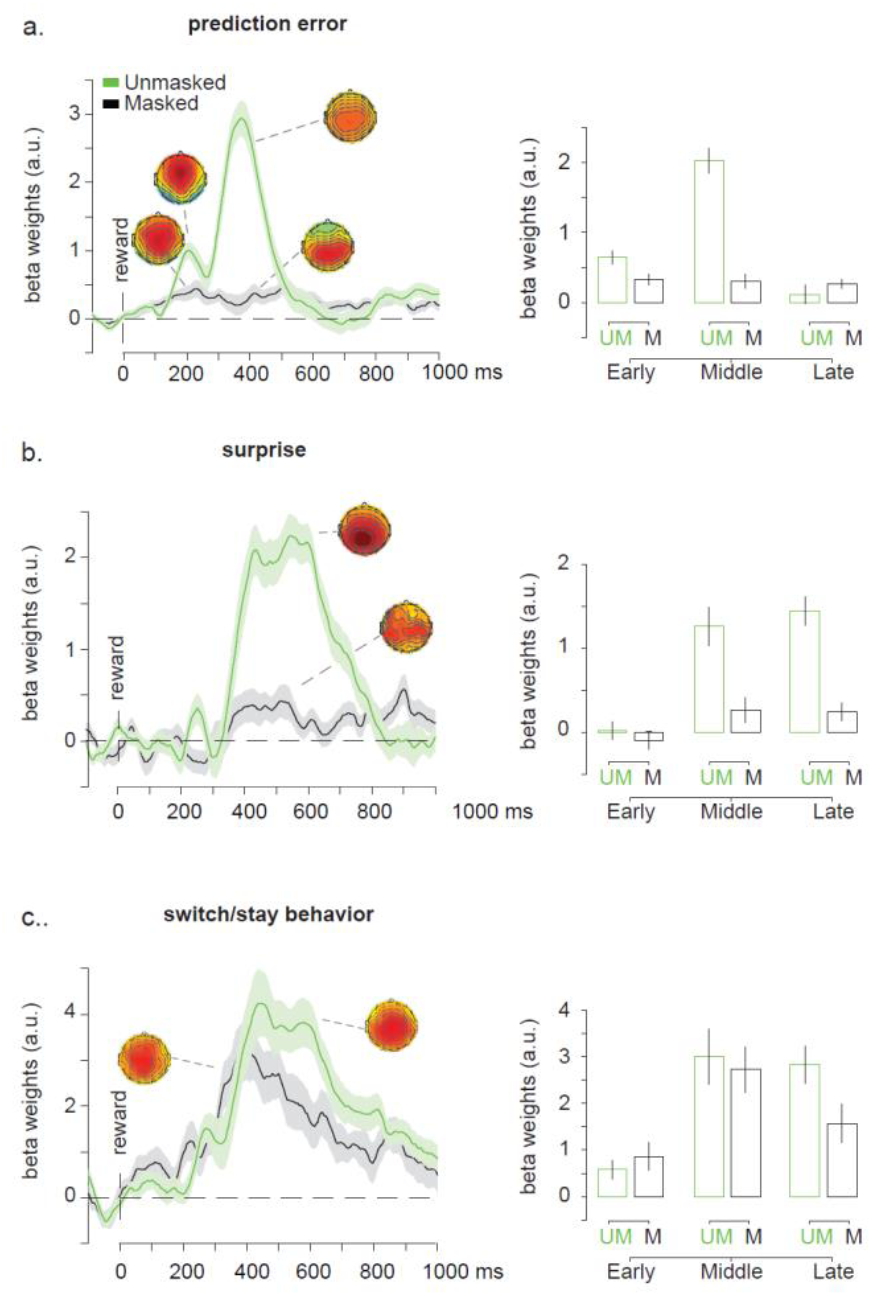
Model-based EEG analysis. **(a)** The time courses of regression weights of the signed PE regressed on the reward-locked EEG signal derived from a central ROI. Effects are plotted separately for unmasked (green) and masked (black) reward outcomes. Shaded areas indicate the s.e.m. Topographical maps show the regression weights during the relevant time windows. Both unmasked and masked reward showed early and mid-latency EEG-PE covariations which are shown in b. Note that the polarities of these components are reversed compared to the ERP results, which in accordance with our expectations, because these ERP modulations are all associated with negative PE values, leading to a reversal of the polarities (maps: 100-300 ms and 300-500 ms; scaling: early masked = [-0.5:0.5], midlatency masked = [-0.5:0.5], early unmasked = [-1:1], middle unmasked = [-3:3]). Bar plots of the signed PE effect for the three time-windows of interest. **(b)** The time courses of regression weights of the unsigned PE, or the level of surprise, regressed on reward-locked EEG signal derived from a central ROI. Both unmasked and masked rewards showed late EEG-surprise covariations (maps: 300-800 ms; scaling: masked=[-0.5:0.5], unmasked=[-2:2]). Bar plots of the surprise effect. **(c)** The time courses of regression weights of switch/stay behavior regressed on the reward-locked EEG signal derived from a central ROI. Both unmasked and masked reward showed late EEG-switch/stay behavior covariations (maps: 300-800 ms; scaling: masked=[-3:3], unmasked=[-3:3]). Bar plots of the switch/stay behavior effect. Error bars represent ± s.e.m. M: masked reward, UM: unmasked reward.

Analyses of the unsigned prediction error signals (i.e. the level of surprise) revealed a rather different pattern of results. For both masked and unmasked reward, and in line with previous findings (Fischer & Ullsperger, 2013; Fouragnan et al., 2017; Mars et al., 2008), this variable was represented in the later P3-like component (time-window 300-500 ms: unmasked: t_31_=5.5, p<0.001; masked: t_31_=1.8, p=0.08; time-window 500-800 ms: unmasked: t_31_=8.4, p<0.001; masked: t_31_=2.2, p=0.03, see **Fig. 5b,** note that headmaps are shown for the middle and late windows combined: 300-800 ms). In both time-windows the effects were stronger for unmasked than masked rewards (all ps<0.001). No significant effects were observed in the early time-window (all ps>0.3).

Finally, we observed a strong relation between switch/stay behavior on the next trial, closely related to the perseveration parameter in the modeling approach, and a broad central positivity (**Fig. 5c**). This effect was already present from the early time-window onwards and was always present irrespective of reward visibility (time-window 100-300 ms: unmasked t_31_=2.9, p=0.006; masked t_31_=2.9, p=0.006; difference t_31_=-0.8, p=0.4; time-window 300-500 ms: unmasked t_31_=5.1, p<0.001; masked t_31_=5.6, p<0.001; difference t_31_=0.5, p=0.6; time-window 500-800 ms: unmasked t_31_=7.1, p<0.001; masked t_31_=3.8, p<0.001; difference t_31_=2.2, p=0.034; **Fig. 5c,** note that headmaps are shown for the middle and late windows combined: 300-800 ms). Interestingly, these effects were very similar for masked and unmasked rewards until ~500 ms after stimulus presentation and significant visibility-related differences only started to emerge in the late time-window. Thus, a larger parietal positive component was associated with an increased likelihood of switching the response option on the next trial. This last analysis not only replicates previous findings about the electrophysiological signature of model-free switching behavior after fully conscious reward (Chase et al., 2011; Fischer & Ullsperger, 2013), but also extends them to the case where reward visibility is very low.

Finally, we ran independent linear regressions with each of the individual EEG-regressor weights shown in the bar plots of **Figure 5** (PE, surprise, switching), for masked and unmasked feedback, for each of the three time-windows of interest, as independent variables and overall performance (percentage correct) as dependent variable, to explore what neural mechanisms correlate with individual performance (18 regressions in total, Bonferoni corrected). Results show that only the middle and late EEG-switching effects from the unmasked feedback (**Fig. 5c**) were positively correlated with performance (both p’s<0.0005).

## Discussion

We combined a reinforcement learning task, a masking procedure, computational modeling and EEG recordings to investigate the impact of reward visibility on different cognitive processes involved in probabilistic reward-guided learning. In behavioral analyses, we observed that participants switched their responses after unmasked and masked unfavorable outcomes (no-reward) more often than after favorable outcomes (reward) (note that masked feedback is not considered “unconscious” here). This pattern of behavior is typically interpreted as evidence for learning. Next, we combined computational modeling with a model-comparison approach. We designed a set of 18 models, built on mixtures of unmasked and masked modules, accounting for reward-based learning and choice perseveration. Reward-based learning was simply operationalized as prediction-error based learning, in line with popular model-free reinforcement-learning algorithms (Berridge, 2004; Dayan & Balleine, 2002; den Ouden et al., 2013; Sutton & Barto, 1998). We then systematically compared the ability of these models to explain our participants’ behavior with a rigorous Bayesian model-comparison approach (Daunizeau et al., 2014). In our model set, which comprised models with and without learning modules from masked feedback, a model including both the masked and unmasked learning modules was identified as the best model. This approach operationalized a clear testing of learning from masked outcomes and provided clear evidence toward the existence of such learning. Our best fitting model also included modules for perseveration after masked and unmasked reward.

An analysis of the best fitting model parameters revealed that learning rates were significantly positive for both visibility modules, although smaller for the masked feedback module. This confirms that participants indeed used both unmasked and masked (although to a lesser extend) reward outcome to inform further decisions. Our results show that the perseveration parameter was also significantly positive for both the visibility modules, although perseveration was smaller for the fully conscious module. This indicates that participants were biased toward repeating previous choices, independently of the outcome of their decisions, an actually frequent observation in human and non-human reinforcement learning tasks (den Ouden et al., 2013; Lau & Glimcher, 2005; Rutledge et al., 2009; Schönberg, Daw, Joel, & O’Doherty, 2007; Seymour et al., 2012). Although often given a low-level interpretation and a connotation of sub-optimality (Voon et al., 2015), perseveration can also constitute the implementation of higher-level behavior: in our task, it is likely that, within a block, participants identified the “good” option based on the integration of information over a long sequence of trials, and therefore decided to ignore irrelevant negative reward basing their choices only on their prior. After masked reward, participants persevered more than after fully conscious reward, revealing that participants stuck to their decision strategy, based on the integration of information over a longer sequence of trials, when full conscious awareness of the outcome was (often) lacking.

Regarding electrophysiological signatures of reinforcement-learning, we observed three neural events evolving over time that were modulated by unmasked outcomes (reward vs no-reward): an early fronto-central FRN, a mid-latency central negativity, and a late centro-parietal P3 component. Crucially, only the fronto-central FRN, which peaked ~200 ms after outcome presentation, was also modulated by masked outcomes. Many studies have reported that this signal, closely related to the response-locked error related negativity (ERN) and originating from the medial frontal cortex (MFC) (Debener et al., 2005; Hauser et. al., 2014), distinguishes positive from negative outcomes (Pfabigan et al. 2011; Cavanagh et al. 2010; Hajcak et al. 2006; Chase et al. 2011; Cohen et al. 2007; Holroyd Nieuwenhuis, Yeung and Cohen, 2003; Fouragnan et al. 2017) in reinforcement-learning tasks (Holroyd & Coles, 2002). This response may reflect a “fast alarm” signal (or alertness response: Fouragnan et al., 2017) that indicates the value of the incoming evidence, which is then accumulated in later stages of the decision making process (Chase et al., 2011; Fouragnan et al., 2017; Ullsperger, Fischer, Nigbur, & Endrass, 2014), possibly reflected in the P3 ERP component (O’Connell, Dockree, & Kelly, 2012). The late parietal P3 ERP component was only observed after fully conscious (unmasked) reward. This signal has been reported to predict behavioral adaptation and the associated update of new stimulus-response associations in memory (Chase et al., 2011; Ullsperger et al., 2014). The P3 has also been related to decision formation and evidence accumulation processes during perceptual decision making (Fischer & Ullsperger, 2013; O’Connell et al., 2012; Ullsperger et al., 2014; Zylberberg, Dehaene, Roelfsema, & Sigman, 2011). Further, our ERP results fit nicely with current theoretical models of conscious and unconscious processes (Dehaene, Charles, King, & Marti, 2014; Lamme, 2006; van Gaal & Lamme, 2012). Within these frameworks, the FRN may reflect a fast feedforward and nonconscious high-level response, whereas the P3 may reflect more conscious, and longer lasting neural responses, potentially dependent on recurrent interactions between distant brain regions (Dehaene & Changeux, 2011).

Although those first EEG analyses outlined important dissociations between learning from reward at different levels of awareness, it is rather difficult to connect these neural signals to precise cognitive processes, using cross trial averaging and traditional contrast-based ERP methods (Cohen & Cavanagh, 2011; Debener et al., 2005; Pernet, Sajda, & Rousselet, 2011; Pfabigan et al., 2011). We therefore ran additional regression analyses in combination with computational modeling to investigate whether single-trial measures of reinforcement learning were influenced by the visibility of probabilistic rewards (Cavanagh et al., 2011; Cohen & Cavanagh, 2011; Pernet et al., 2011). We focused our investigations on the EEG-correlates of three main computational variables: the prediction-error (signed PE), the level of surprise (unsigned PE) and switch/stay behavior on the next trial. This analysis revealed a striking similarity of neural PE correlates after both unmasked and masked reward outcomes, although weaker for the latter. Both the early and mid-latency negative ERP components were associated with PE computation (Fouragnan et al. 2017), whereas the parietal P3 was not. These findings support previous results showing that the FRN reflects signed PE signals (Holroyd & Coles, 2002; Overbeek, Nieuwenhuis, & Ridderinkhof, 2005), likely emerging from dopaminergic projections to the MFC (Jocham, Klein, & Ullsperger, 2011; Park et al., 2012; Schultz, 2007; Walsh & Anderson, 2012), although especially the early response has also been linked to noradrenergic and serotonergic modulations (discussed in Fouragnan et al. 2015).

Interestingly, whereas the two early neural events coded for a signed PE signal, the later P3 component was particularly modulated by the unsigned PE, reflecting the level of surprise. Although this corroborates similar results obtained with different techniques and methods (Fouragnan et al., 2017; Mars et al., 2008), we crucially show here that the level of surprise is also encoded in parietal EEG fluctuations elicited by masked reward outcomes. Finally, the EEG-switch/repeat correlations that we report here are in line with previous studies showing that trial-by-trial switch behavior can be observed at parietal channels as a late positive P3 component (Chase et al., 2011; Fischer & Ullsperger, 2013). In a previous study in which the authors combined computational modeling and RL learning it has been shown that this neural event did not differ when participants received actual reward about their choice or merely fictive reward (Fischer & Ullsperger, 2013). Here we show that this effect likely represents decision strategies that are formed over longer timescales. Overall, these results show that several cognitive processes important for reward-based learning, namely PE computation, surprise and switch/stay implementation are processed in the human brain and these cognitive processes are temporally and spatially dissociated in time (Fouragnan et al. 2017).

### Future directions, open questions and limitations

Although several crucial question about the role of feedback awareness in reward-based learning were addressed here, several interesting questions remain unanswered. First, the current task design did not allow us to analyze what neural processes may drive “correct switching behavior” vs. switching behavior in general, due to the low number of block reversals and therefore low number of possible correct switch trials (max. 11 trials per subject). Future studies may address this issue by incorporating more volatile reward environments, containing more block reversals (and therefore correct switches), to address this issue (see e.g. Behrens, Woolrich, Walton, & Rushworth, 2007). Another open question relates to the isolation of the neural and cognitive processes underlying the early vs. mid-latency frontal ERP negativities. Previous studies have typically observed only one frontal negativity (the FRN), instead of two (for reviews see Cavanagh & Frank, 2014; Cohen et al., 2011). At present, it remains unclear why this is the case and future work is necessary to unravel the task specifics that may drive these differences between studies. The combination of both EEG and fMRI, as done previously (Debener, Ullsperger, Siegel, & Engel, 2006; Fouragnan et al., 2017; Hauser et al., 2014) may contribute to this endeavor. Finally, future studies are crucial to explore what factors may drive that the model-based single-trial regressions yielded weaker (but often still significant) effects for the masked condition compared to the unmasked condition. An interesting option may be that on a subset of trials masked feedback could have been completely missed by the system, such that no prediction error could be generated (and represented in the EEG).

## Author contributions

SVG and CMCC conceived the experiment. CMCC, SN and JJ performed the experiments. CMCC, SN and SVG performed EEG analyses. ML and SP performed modeling analyses. CMCC, SP, ML and SVG wrote the manuscript. MXC and ML provided expertise and feedback. SVG supervised the project.

